# Glutathione *S*-transferases: unexpected roles in astrocyte activation and astrocyte-microglia communication during brain inflammation

**DOI:** 10.1101/199612

**Authors:** Shin-ichi Kano, Eric Y. Choi, Eisuke Dohi, Indigo V. L. Rose, Daniel J. Chang, Ashley M. Wilson, Brian D. Lo, Takashi Imai, Akira Sawa

## Abstract

Astrocytes and microglia play critical roles in brain inflammation, but their mutual regulation is not fully understood. Here we report unexpected roles for glutathione *S*-transferases (GSTs), particularly GSTM1, in astrocyte activation and astrocyte-mediated enhancement of microglia activation during brain inflammation. We found that astrocyte-specific silencing of GSTM1 expression in the prefrontal cortex (PFC) attenuated microglia activation in brain inflammation induced by systemic injection of lipopolysaccharides (LPS). *Gstm1* silencing in astrocytes also attenuated LPS-induced TNF-α production by microglia in co-culture. In astrocytes, GSTM1 was required for the activation of nuclear factor-κB (NF-κB) and c-Jun N-terminal kinases (JNK) and the production of pro-inflammatory mediators previously implicated in microglia activation, such as granulocyte-macrophage colony-stimulating factor (GM-CSF/CSF2) and chemokine (C-C motif) ligand 2 (CCL2). Similar results were also obtained with GSTT2 both *in vitro* and *in vivo*. Thus, our study identified a critical role for GSTs in priming astrocytes and enhancing microglia activation during brain inflammation.

**Significant Statement:** Astrocytes and microglia play critical roles in brain inflammation, but it is not fully understood how astrocytes regulate microglia activation. Here we report a novel mechanism by which glutathione *S*-transferases (GSTs), the enzymes for phase II detoxification of xenobiotic metabolism, in astrocytes control microglia activation during brain inflammation. We found that GSTs, particularly GSTM1, regulate the induction of pro-inflammatory mediators via the activation of NF-κB and JNK in astrocytes. Our studies provide evidence that GST enzymes are active players in brain inflammation and can be targeted to regulate microglia activation.

## Introduction

Astrocytes play a critical role in maintaining normal neuronal function by modulating synaptic activity, supporting neuronal survival, and providing metabolic support (1-4). In brain inflammation, astrocytes have been suggested to regulate the activity of microglia, neurons, oligodendrocytes, and immune cells infiltrating from the periphery (4-6). Because both astrocytes and microglia sense immune stimuli and produce inflammatory mediators, it is important to understand the mechanisms by which astrocytes and microglia regulate each other. Nonetheless, it is not fully understood how astrocytes influence microglia activation in brain inflammation.

Glutathione (GSH) is a thiol-containing tripeptide and a major antioxidant within cells (7). Decreases in GSH and increases in the oxidized form of GSH, GSSG, are associated with cellular susceptibility to oxidative stress. GSH also influences cellular functions through *S*-glutathionylation, the reversible conjugation of a GSH molecule to reactive cysteine residues in proteins (8, 9). Dysregulation of glutathione metabolism is associated with brain inflammation in various neurological and psychiatric disorders (10-17). It is not clear, however, how the glutathione system influences inflammatory responses at the mechanistic level.

Glutathione *S*-transferases (GSTs) are the enzymes conjugating the reduced glutathione (GSH) to target molecules in the phase II detoxification of drug metabolism (18). GSTs consist of a diverse family of cytosolic, mitochondrial, and microsomal enzymes and prevent cellular damage from noxious stimuli of xenobiotic metabolites (18-20). GSTs are widely expressed throughout the body with abundant expression in the liver, kidney, and lung (18-20). In addition, some of the mu, pi, and alpha classes of GSTs (GSTM1, GSTP1 and GSTA4) were detected in the human and rodent brains (21-23). Notably, a recent study suggested that GSTM1 is one of the most abundantly expressed proteins in astrocytes (24).

Accumulating evidence shows that GSTs also influence a wide range of biological mechanisms, such as redox homeostasis, signal transduction, cell proliferation, and cell death (25-27). GSTs are known to exert these regulatory functions by activating/inhibiting their target molecules, such as c-Jun N-terminal kinases (JNK), apoptosis signal-regulating kinase 1 (ASK1), and nuclear factor-κB (NF-κB) through either protein-protein interactions or *S*-glutathionylation (27, 28). The current knowledge about these non-phase II detoxification roles of GSTs, however, is still very limited. In the brain, neuronal GSTP1 has been shown to protect neuronal cell death in an animal model of 1-methyl-4-phenyl-1,2,3,6-tetrahydropyridine (MPTP)-induced degeneration of dopaminergic neurons in the substantia nigra (29). In contrast, the roles of specific GSTs in glial cells have not been well characterized, particularly *in vivo*.

Genetic studies also suggest that variations in the genes encoding GSTs are involved in neurological and psychiatric disorders with immune dysregulation, such as Parkinson’s disease, Alzheimer’s disease, multiple sclerosis, schizophrenia, and autism (16, 30-34). More recently, altered expression of GST activity-related genes has been found to be one of the most drastically changed molecular signatures, together with immune and microglia-related genes, in postmortem brains from patients with late-onset Alzheimer’s disease (35). Nevertheless, it is not known whether GSTs are involved in the regulation of astrocytes and microglia during brain inflammation.

Here we investigated the role of GST enzymes in astrocytes in a mouse model of brain inflammation. We found that GSTs in astrocytes, particularly GSTM1 and GSTT2, are required for the activation of microglia in brain inflammation induced by systemic LPS administration. Mechanistically, GSTM1 was shown to activate NF-κB and JNK and enhance the induction of both pro-inflammatory mediators such as granulocyte-macrophage colony-stimulating factor (GM-CSF/CSF2) and chemokine (C-C motif) ligand 2 (CCL2). Thus, we propose that GSTs prime the inflammatory responses of astrocytes and enhance the activation of microglia via secretion of pro-inflammatory mediators.

## Results

### Enriched expression of GSTM1 and GSTT2 in astrocytes in the mouse brain

While previous studies showed that GSTM1 and GSTP1 proteins were expressed in the mouse brain (21, 29), region and cell-type specificity of their expression patterns was not fully addressed. In addition, it was not clear whether other GST enzymes, such as GST theta (GSTT), were also expressed as proteins in the brain. A recent study reported that GSTM1 is one of the most abundantly expressed proteins in astrocytes (24). Thus, we examined the expression patterns of GSTM1 and GSTT2 in various regions of the mouse brain. As expected, GSTM1 was abundantly expressed in the mouse brain, including the cortex, hippocampus, striatum, and cerebellum (**Fig. 1A**). The expression pattern of GSTT2 also showed a similar pattern although its expression level was much lower. Immunohistochemical analysis revealed that both GSTM1 and GSTT2 were enriched in astrocytes compared to neurons, oligodendrocytes, and microglia (**Fig**. **1B** and **Fig. S1**). These results suggest that GSTM1 and GSTT2 primarily regulate astrocyte function in the mouse brain.

**Fig. 1.**
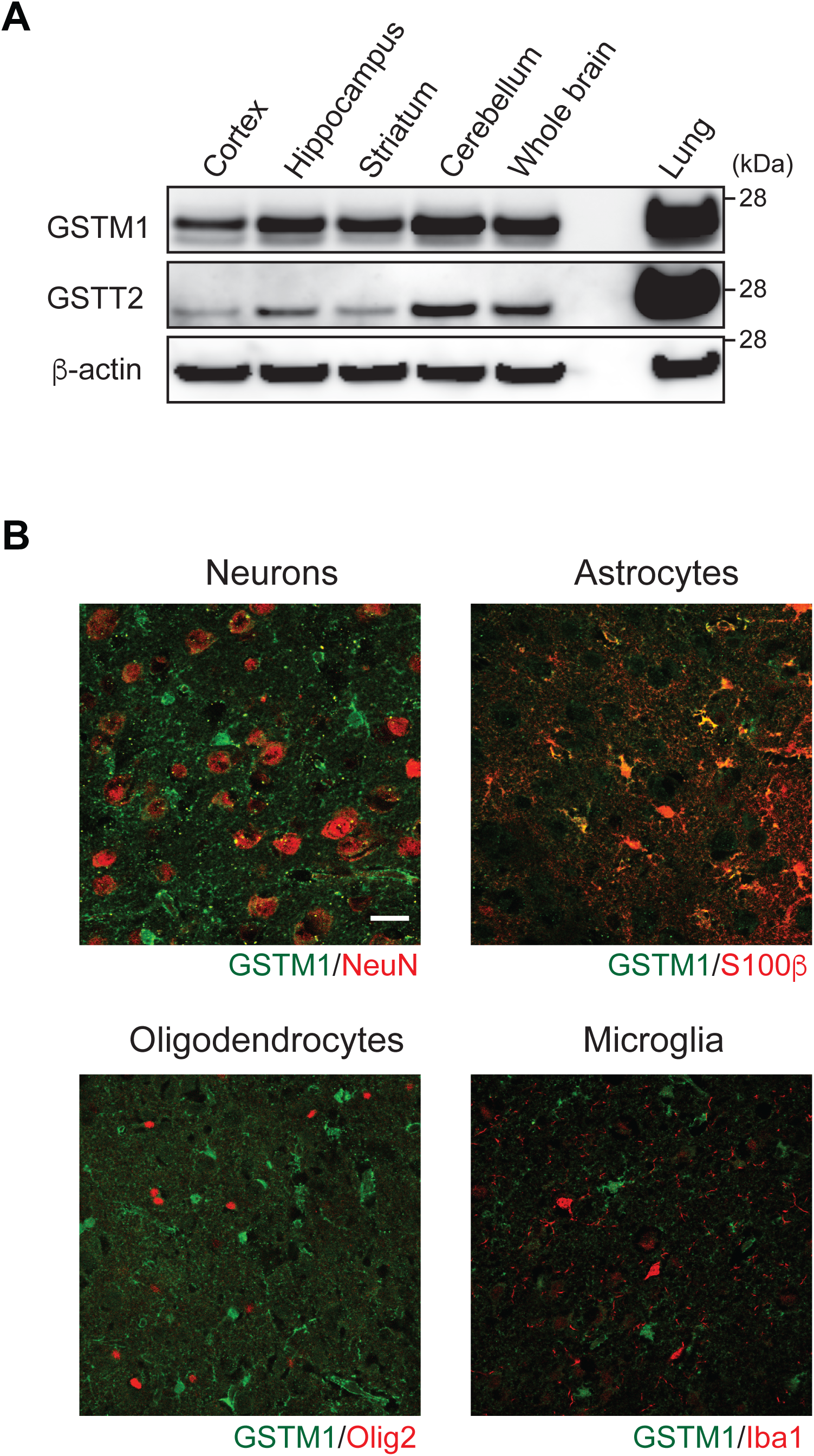
Enriched expression of GSTM1 in astrocytes in the mouse brain. **A.** Expression of GSTM1 and GSTT2 in the cerebral cortex, hippocampus, striatum, and cerebellum of C57BL/6 wild-type mice (8 weeks of age) by Western blot. **B.** Co-localization of GSTM1 in astrocytes in the medial prefrontal cortex (mPFC), specifically prelimbic area (PrL). Immunofluorescent staining of GSTM1 protein in the cerebral cortex of 8 weeks old C57BL/6J mice was performed using GSTM1 antibody together with cell-type specific markers (NeuN for neurons, S100β for astrocytes, Olig2 for oligodendrocytes, and Iba1 for microglia). Scale bar, 25 μm.

### Requirement of GSTM1 and GSTT2 in astrocytes for microglia activation *in vivo*

Because astrocytes play a critical role in brain inflammation, we examined the impact of GSTM1 and GSTT2 silencing in astrocytes on inflammatory responses in the brain. Systemic administration of lipopolysaccharides (LPS) is a well-established model to induce brain inflammation, and is characterized by the activation of microglia and astrocytes (36, 37). We utilized this model to examine the effect of astrocyte-specific knockdown of GSTs on the activation of microglia. We first knocked down GSTM1 expression by using an adeno-associated virus (AAV) vector encoding green fluorescent protein (GFP) and mir-30-based short hairpin RNA targeting *Gstm1* (*Gstm1* shRNAmir) downstream of a floxed stop codon (AAV-LSL-GFP-*Gstm1*-shRNAmir) (**Fig. S2**). The AAV was stereotactically injected into the medial prefrontal cortex (mPFC) of mice expressing Cre recombinase under the mouse *Gfap* gene promoter (m*Gfap*-Cre mice). As shown in **Fig. 2B**, the expression of GFP was specific for astrocytes (S100β^+^ cells) in the mPFC of m*Gfap*-Cre mice at 3 weeks after the AAV injection. No GFP signals were observed in neurons (NeuN^+^ cells), oligodendrocytes (NG2^+^ cells), or microglia (Iba1^+^ cells) (data not shown). Under this condition, we injected LPS intraperitoneally and examined the activation status of microglia in the area where GFP^+^ astrocytes were detected. An AAV virus carrying a non-targeting (NS) shRNAmir was used as a control (control shRNAmir). We found that *Gstm1* silencing in astrocytes attenuated the activation of nearby microglia, judged by their morphological changes, at 48 h after LPS injection (**Fig. 2C** and **Fig. S3**). The expression of TNF-α was also diminished in these microglia (**Fig. 2D**). Microglia activation was similarly attenuated when GSTT2 expression was silenced in astrocytes by injection of AAV-LSL-GFP-*Gstt2*-shRNAmir virus into m*Gfap*-Cre mice (**Fig. S4**). These data showed that GSTM1 and GSTT2 in astrocytes are required for the activation of microglia during brain inflammation.

**Fig. 2.**
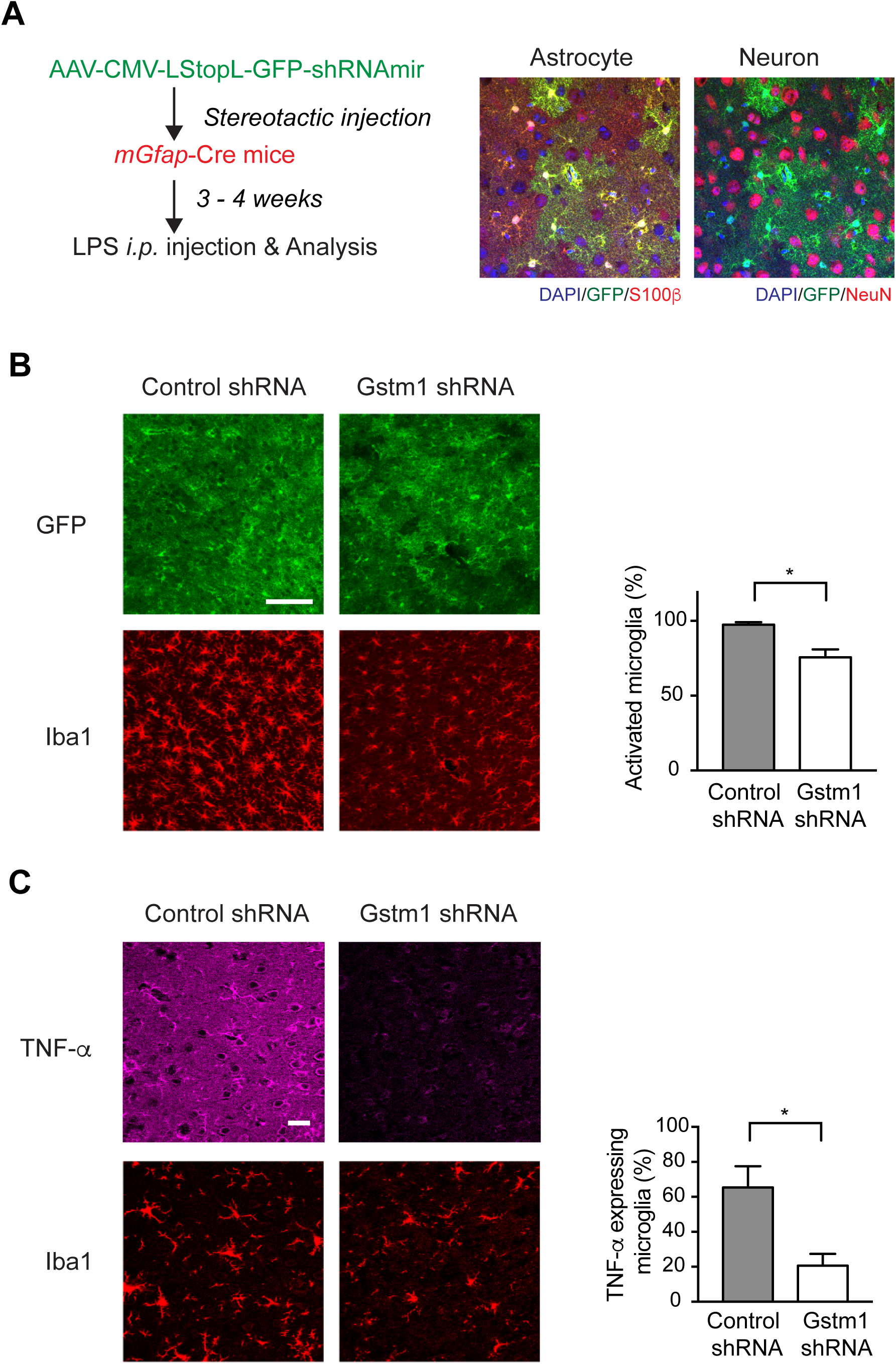
Reduced activation of microglia in the mice with GSTM1 knockdown in astrocytes under brain inflammation induced by systemic injection of LPS. **A.** Experimental design. Mouse *Gfap* gene promoter-driven Cre transgenic (m*Gfap*-Cre) mice were stereotactically injected with floxed-AAV vector encoding shRNAmir against *Gstm1* (AAV-LSL-GFP-*Gstm1* shRNAmir) into the medial prefrontal cortex (mPFC) and challenged with intraperitoneal injection of LPS (5 mg/kg) 3-4 weeks later. After 48 h, the brains were harvested and stained for the presence of virally encoded GFP together with cell-type specific markers (NeuN for neurons and S100β for astrocytes). **B.** Reduced activation of microglia in the vicinity of astrocytes with GSTM1 knockdown (GFP^+^). Microglia were stained with a marker, Iba1, and their activation status was estimated by their morphological changes. To characterize microglial activation, we distinguished each Iba1^+^ microglia based on their morphology either ramified, intermediate, amoeboid, or round (53). Only microglia that show ramified morphology was counted as resting and those that showed intermediate, amoeboid, or round morphology was counted as activated (**Fig. S3**). Graph shows the percentage of activated Iba1^+^ microglia per total Iba1^+^ microglia from the mice injected with control shRNA (9 slices; 3 mice) and Gstm1 shRNA (9 slices; 3 mice). **C.** Decreased levels of TNF-α expression in microglia in the vicinity of astrocytes with GSTM1 knockdown. Graph shows percentage of TNF-β expressing Iba1^+^ microglia per total Iba1^+^ microglia. Scale bars, 100 μm (**B**) and 25 μm (**C**). Each bar represents mean ± s.e.m. Significance was determined by Student’s *t*-test. \**p*<0.05.

### Non-cell autonomous effect of GSTM1 and GSTT2 in astrocytes on microglia activation in co-culture

Microglia and astrocytes can amplify each other’s activation by secreting pro-inflammatory mediators (**Fig. 3A**)(38). To understand the mechanisms underlying the effects of GSTM1 knockdown in astrocytes on the activation of microglia, we utilized a co-culture system of primary mouse astrocytes and BV2 microglia (**Fig. 3B)**. Purified primary mouse cortical astrocytes (**Fig. S5**) were infected with a lentivirus encoding shRNA targeting *Gstm1* (*Gstm1* shRNA) or control shRNA, and then mixed with BV2 microglia. In this co-culture system, TNF-α production is totally dependent on the presence of microglia (**Fig. 3B**). We then compared the effects of *Gstm1* silencing in astrocytes on microglial TNF-α production. Consistent with our *in vivo* findings, *Gstm1* silencing in astrocytes reduced the amount of TNF-α secretion and its mRNA induction after LPS stimulation (**Fig. 3C** and **D**). Notably, inhibition of TNF-α and IL-1α signaling by blocking antibodies attenuated the mRNA induction of TNF-α, GM-CSF (CSF2), and CCL2 (**Fig. 3E**). These data support the presence of active communication between astrocytes and microglia, and that GSTM1 in astrocytes is required for microglial TNF-α production in a non-cell autonomous manner. GSTM1 or GSTT2 overexpression in astrocytes, on the other hand, enhanced induction of TNF-α mRNA in co-culture (**Fig. S6**).

**Fig. 3.**
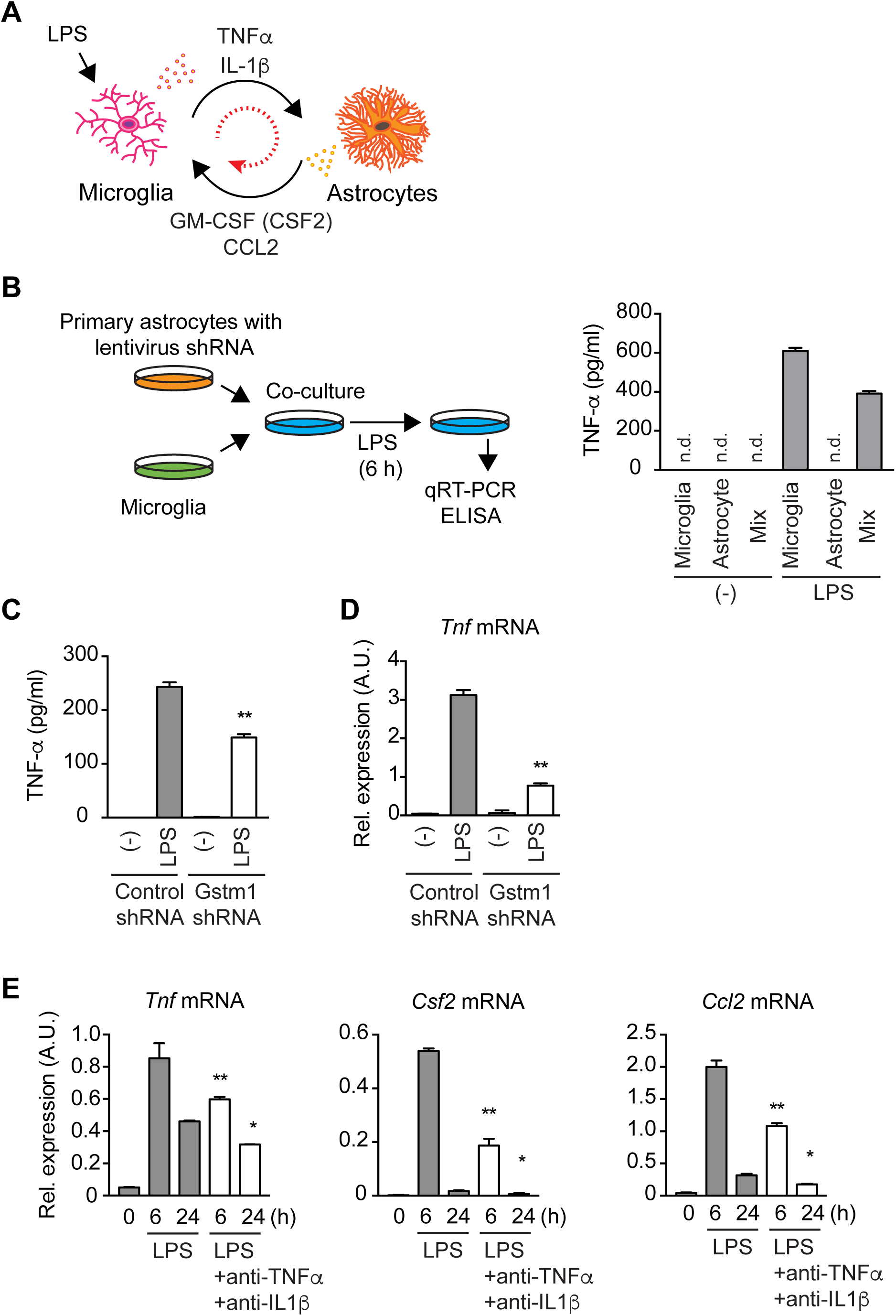
Impaired production of microglial TNF-α by GSTM1 silencing in co-cultured astrocytes. **A.** Amplification of inflammatory responses between astrocytes and microglia via soluble mediators. Previous studies suggest that microglia produce pro-inflammatory cytokines such as TNF-α and IL-1β, which in turn stimulate astrocytes to produce pro-inflammatory mediators such as GM-CSF (CSF2) and CCL2. **B.** Experimental design. Primary mouse cortical astrocytes were infected with lentivirus encoding shRNA, and then co-cultured with BV2 microglia. Co-cultures were stimulated with LPS (1 μg/ml) for 6 h. Cells and culture supernatants were harvested for qRT-PCR analysis and ELISA, respectively. As shown in the graph, microglia, but not astrocytes, produce TNF-α in response to LPS stimulation in this co-culture. **C.** Reduced TNF-α production in the co-culture of BV2 microglia and astrocytes with GSTM1 silencing by ELISA. **D.** Reduced *Tnf* mRNA expression in the co-culture of BV2 microglia and astrocytes with GSTM1 knockdown by qRT-PCR analysis. **E.** Reduced mRNA expression for *Tnf*, *Csf2*, and *Ccl2* genes in co-culture of wild-type astrocytes and BV2 microglia after blocking TNF-α and IL-1α signaling. Each bar represents mean ± s.e.m.; n.d., not detected. For **B**, representative data from two independent experiments were shown, and for **C**-**E**, representative data from three independent experiments were shown. Significance was determined by two-way ANOVA with Sidak’s post-hoc test. \**p*<0.05, \*\**p*<0.01

### Control of the production of inflammatory mediators by GSTM1 in astrocytes

To investigate the molecular basis on which GSTM1 regulates astrocyte function during inflammatory responses, we analyzed the effects of *Gstm1* silencing on purified primary mouse astrocytes. Previous studies showed that astrocytes produce GM-CSF (CSF2) and CCL2, both of which are potent activators of microglia (38-41), during brain inflammation. Thus, we examined their secretion from astrocytes in response to pro-inflammatory cytokines TNF-β and IL-1β. We found that the amount of GM-CSF (CSF2) and CCL2 in the culture supernatants from astrocytes decreased in the absence of GSTM1 (**Fig. 4A**). We also performed qRT-PCR analysis to examine mRNA induction of pro- and anti-inflammatory mediator genes. The induction of *Ccl2, Csf1, and Csf2* mRNA, but not *Nos2* mRNA, was impaired by *Gstm1* silencing in astrocytes (**Fig. 4B**). *Gstm1* silencing also reduced the level of *Tgfb1* mRNA irrespective of the presence of TNF-β and IL-1β. In contrast, mRNA induction of *Il33*, another immunoregulatory cytokine, was enhanced in the absence of GSTM1 (**Fig. 4B**). These findings showed that GSTM1 regulates the transcription of both pro- and anti-inflammatory genes in astrocytes.

**Fig. 4.**
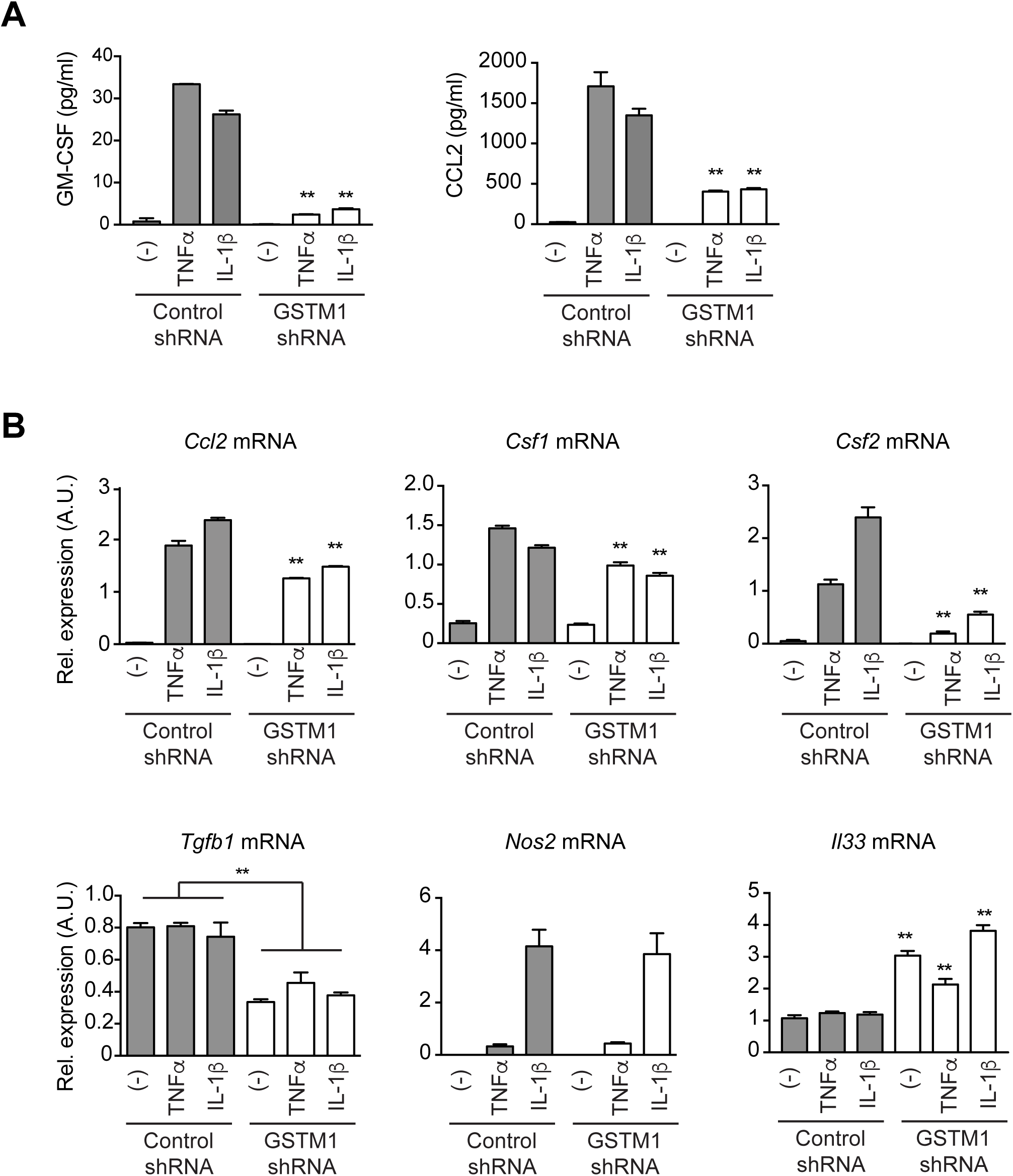
Altered induction of inflammatory mediators in astrocytes with GSTM1 knockdown. **A.** Reduced secretion of GM-CSF (CSF2) and CCL2 by astrocytes with GSTM1 knockdown in response to TNF-β and IL-1β. Culture supernatants from mouse cortical astrocytes stimulated with TNF-β (50 ng/ml) and IL-1β (10 ng/ml) for 6 h were analyzed by ELISA. **B.** Expression of *Ccl2*, *Csf1*, *Csf2*, *Tgfb1*, *Nos2,* and *Il33* mRNAs in primary mouse cortical astrocytes with GSTM1 and control knockdown. Primary mouse cortical astrocytes were stimulated with TNF-β (50 ng/ml) and IL-1β (10 ng/ml) for 6 h and their mRNA expression was examined by qRT-PCR analysis. Each bar represents mean ± s.e.m. Representative data from at least two independent experiments were shown. Significance was determined by two-way ANOVA with Sidak’s post-hoc test. ** *p*<0.01.

### Mechanisms of GSTM1-mediated activation of TNF-αand IL-1β signaling in astrocytes

To address the mechanisms underlying altered gene transcription downstream of TNF-α and IL-1β in astrocytes, we analyzed the activation of NF-κB, ERK, and JNK cascades in primary mouse astrocytes. As shown in **Fig. 5A**, *Gstm1* silencing in astrocytes reduced phosphorylation levels of NF-κB p65 and JNK in response to either TNF-α or IL-1β. ERK phosphorylation was reduced by *Gstm1* silencing, but not induced by either TNF-α or IL-1β stimulation. We further found that *Gstm1* silencing attenuated the phosphorylation of IκBα (**Fig. 5B**). IκBα phosphorylation is a rate limiting step for the translocation of NF-κB p65/p50 components from the cytoplasm into the nucleus to activate their target genes (42, 43). Thus, GSTM1 controls NF-κB signaling upstream of IκBα phosphorylation, possibly at the level of the activation of IκB kinases (IKKα, IKKβ, and IKKγ). Because previous studies reported that pharmacological depletions of GSH dampened TNF-α-induced activation of the NF-κB pathway in primary mouse hepatocytes (44), we next examined the effects of GSH depletion by diethylmaleate (DEM) on TNF-β-induced phosphorylation of p65 in mouse astrocytes. Notably, similar to *Gstm1* silencing, GSH depletion caused reduced levels of phosphorylation of p65 in response to TNF-α (**Fig. 5C**). Thus, these findings indicate that GSTM1 activates NF-κB and JNK signaling and induces the expression of pro-inflammatory mediators such as GM-CSF (CSF2) and CCL2, which enhance the activation of microglia (**Fig. 5D**).

**Fig. 5.**
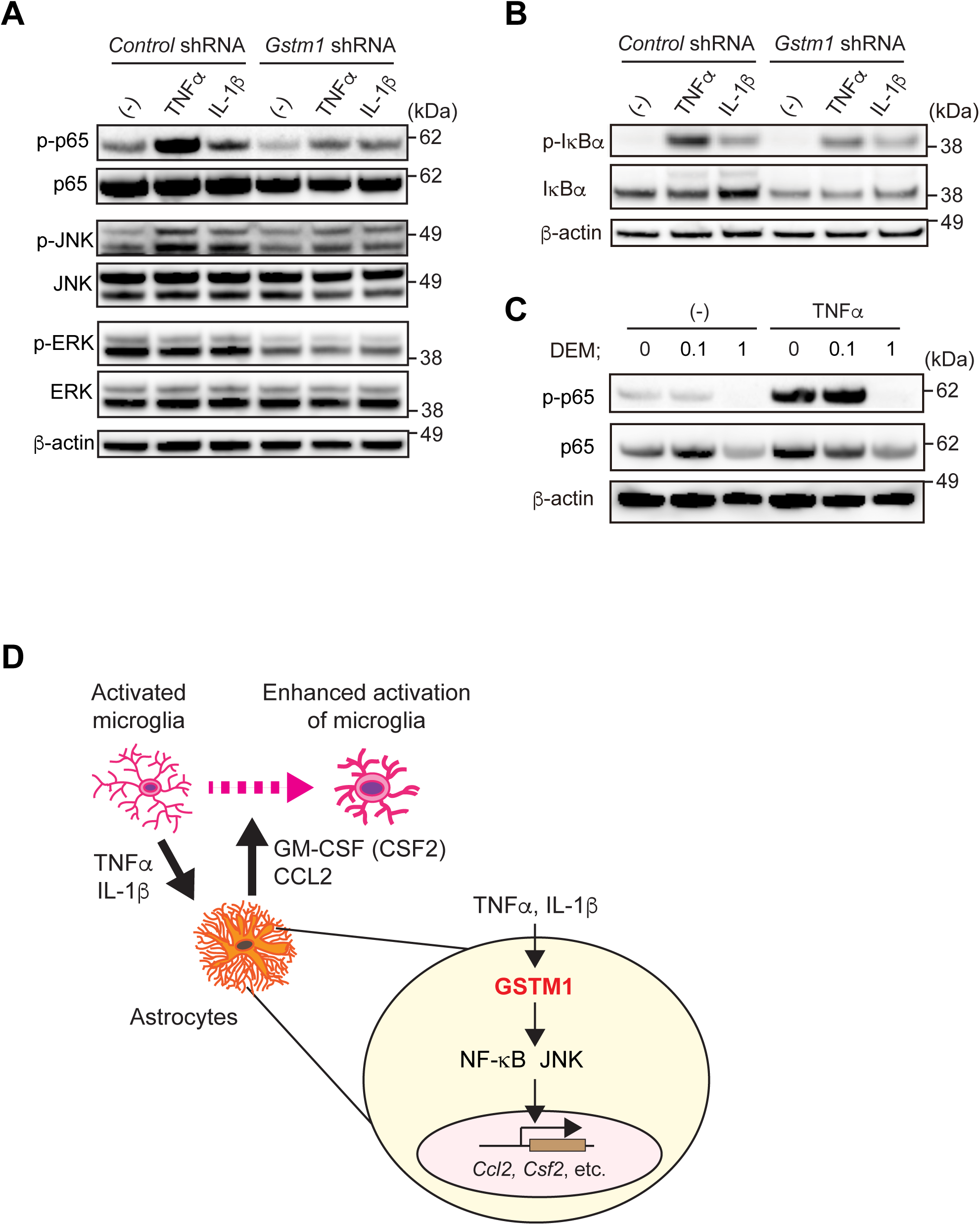
Activation of NF-κB and JNK cascades by GSTM1 in purified astrocytes. **A.** Astrocytes with GSTM1 and control knockdown were stimulated with TNF-β (50 ng/ml) and IL-1β (10 ng/ml) for 6 h and the phosphorylation and expression levels of NF-κB p65 subunit, JNK, and ERK, together with the expression of β-actin, were examined by Western blot. **B.** Decreased phosphorylation of IκB-β in astrocytes with GSTM1 knockdown. **C.** Decreased activation of NF-κB p65 in astrocytes by DEM treatment (DEM 0.1 mM and 1 mM) for 6 h during TNF-β (50 ng/ml) stimulation. **D**. A schematic model on the role of GSTM1 in astrocytes and astrocyte-microglia interaction. Our findings support the role of GSTM1 in activating NF-κB and JNK and inducing the expression of *Ccl2* and *Csf2* genes in astrocytes. Thus, in the absence of GSTM1, microglia activation is attenuated by insufficient amount of astrocyte-derived GM-CSF (CSF2) and CCL2. For **A**-**C**, representative data from at least two independent experiments were shown.

## Discussion

In this study, we discovered that two GST enzymes, GSTM1 and GSTT2, in astrocytes are required for the activation of microglia during brain inflammation induced by systemic LPS administration. In addition, GSTM1 was shown to regulate the induction of pro- and anti-inflammatory mediators in astrocytes such as GM-CSF (CSF2) and CCL2. We demonstrated that GSTM1 in astrocytes modified NF-κB and JNK signaling pathways, possibly via protein *S*-glutathionylation. These data have revealed a critical role of GST enzymes in astrocytes by enhancing the activation of microglia in brain inflammation.

Astrocytes are widely involved in brain inflammatory responses related to infection, autoimmunity, neurodegeneration, injury, and other pathological conditions (1-3, 5, 6). While astrocyte-neuron interactions have been extensively studied, less is known about the interactions between astrocytes and microglia in brain inflammation. Because both of these glial cells sense immune stimuli and produce inflammatory mediators, it is important to understand their mutual regulation. Our data suggest that astrocytes control the activation of microglia, at least in LPS-induced brain inflammation. Recent studies have highlighted the role of microglia in the functional polarization of reactive astrocytes, such as A1/A2 astrocytes, in the same model of brain inflammation (4, 36, 37). Thus, it is speculated that the interaction of microglia and astrocytes forms a feed-forward loop to facilitate inflammatory responses in the brain. Further studies will reveal whether this GST-mediated regulatory mechanism is common to other brain inflammation models, such as infections, autoimmune diseases, and neurodegeneration. In addition, the impact of GSTs on the functions of astrocytes themselves needs to be determined.

Although GSTP was previously studied in an animal model of 1-methyl-4-phenyl-1,2,3,6-tetrahydropyridine (MPTP)-induced degeneration of dopaminergic neurons in the substantia nigra (29), it was not clear whether GST enzymes are involved in more physiological inflammatory conditions beyond xenobiotics-induced neurodegeneration. Our study demonstrated that GSTM1 and GSTT2 are required for LPS-induced brain inflammation unrelated to xenobiotics such as MPTP, adding to the increasing body of work that support the role of GSTs as endogenous regulators of physiological processes distinct from phase II detoxification of drug metabolism (25-27). It is speculated that many proteins are *S*-glutathionylated by GSTs in various mouse and human cells (25-27). The identification of those target proteins will further facilitate our understanding of immunoregulatory mechanisms by diverse GSTs in different tissues and cell types.

Based on our findings, we propose that GST variations in human populations may dictate individual variations in inflammatory responses or susceptibility to immune pathology associated with various acute and chronic diseases. Despite numerous genome-wide association studies and exome sequencing studies published so far, there has been no clear strong genetic evidence supporting the role of GST variations in specific diseases. This indicates that GST variations are not associated with specific disease(s) but rather modify individual responses to homeostatic imbalance, such as inflammation, in a wide range of disorders. As GST enzymes belonging to different classes are distinct in protein structures (18-20), small molecule inhibitors/activators for GST enzymes may be useful as add-on therapeutic agents to modify the outcome of inflammation caused by various diseases.

## Materials and Methods

### Mice

*mGfap*^*Cre*^ *mice* (lines 73.12 and 77.6) and C57BL/6J mice were purchased from the Jackson Laboratory. C57BL/6 timed pregnant female mice for *in vitro* cell cultures were purchased from Charles River Laboratories. Mice were housed in specific pathogen free facilities at the Johns Hopkins University. All the procedures were approved by the Institutional Animal Care and Use Committee of the Johns Hopkins University.

### Virus preparation

Lentiviruses (pLKO.1 and pGIPZ viruses) and adeno-associated viruses (AAVs) were prepared following our established protocols (45-47) and their titers were estimated by quantitative PCR (qPCR)-based methods (45, 46, 48). For lentiviruses, pLKO.1-*Gstm1* shRNA lentiviral vectors (#1, TRCN0000103241; #2, TRCN0000103243; #3, TRCN0000103244; #4, TRCN0000103240) were obtained from the RNA consortium (TRC) library via the Hit center at the Johns Hopkins University School of Medicine; control pLKO.1-*Gstm1* shRNA lentiviral vector (#30323) was obtained from Addgene; and pGIPZ-*Gstt2* shRNAmir lentiviral vectors (#1, V2LMM_67055; #2, V2LMM_218573; #3, V3LMM_449685; #4, V3LMM_449688) and control non-silencing (NS) shRNAmir lentiviral vector (RHS4346) were obtained from the Open Biosystems. pHAGE-Gstm1 and pHAGE-Gstt2 were generated by subcloning mouse Gstm1 and Gstt2 cDNAs into pHAGE vectors. For AAVs, AAV-loxP-Stop-loxP (LSL)-GFP-*Gstm1* shRNAmir was generated based on the most efficient shRNA construct (#4) (**Fig. S2**) following the established protocol (49); and AAV-LSL-GFP-*Gstt2* shRNAmir and AAV-LSL-GFP-NS shRNAmir were generated by shuttling *Gstt2*-shRNAmir (#2) and NS-shRNAmir from pGIPZ lentiviral vectors into AAV-LSL-GFP vectors.

### Cell culture

Primary mouse glial cell cultures were prepared from the cortices of postnatal day 3 (P3) pups of C57BL/6 mice as described previously (50-52). After careful removal of meninges, single cell suspensions were obtained by serial trituration of cerebral cortices with 20G and 26G needles with a 10-ml syringe, followed by the removal of cell debris with a 40-μm cell strainer (Fisher Scientific). Cells were suspended in DMEM/F12 supplemented with 15% FBS and penicillin/streptomycin (all from Thermo Fisher Scientific), and seeded onto T-75 flasks (Corning) pre-coated with poly-D-lysine (PDL, 25 μg/ml) at approximately two brains per flask. Medium was changed on day 3, and every other day thereafter. If necessary, lentiviral infection was performed on day 10 as described below. To enrich astrocytes, oligodendrocyte lineage cells and microglia on the surface of mixed glial cell culture were vigorously shaken off and astrocytes were then collected as negative fractions after MACS sorting with CD11b Microbeads (Miltenyi Biotec). Collected astrocytes were >98% GFAP^+^ CD11b^+^ cells. BV2 microglia were maintained in DMEM/F12 supplemented with 15% FBS and penicillin/streptomycin (all from Thermo Fisher Scientific). Astrocyte-microglia co-cultures were prepared by seeding 5 x 10^5^ astrocytes onto PDL-coated 6-well plates and adding 5 x 10^4^ BV2 cells 2 days later.

### Lentiviral infection

Lentiviruses were added to primary glial cell cultures on day 10 at 1:1 MOI (multiplicity of infection). For pLKO.1 lentiviruses, the virus-infected cells were enriched by antibiotics-based selection (puromycin, 2.5 μg/ml) for 72 h beginning at 72 h post-infection. For pHAGE lentiviruses, infection efficiency was monitored by GFP signals under a fluorescence microscope.

### *In vitro* cell stimulation

Astrocyte-microglia co-cultures and purified astrocytes were stimulated with 1 μg/ml of LPS (O55B5, Sigma) or cytokines [TNF-α (50 ng/ml) and IL-1β (10 ng/ml)] for 6 h, respectively. For GSH depletion experiments using diethylmaleate (DEM, Santa Cruz Biotechnology), purified astrocytes were seeded with a density of 5 x 10^5^ cells per well onto PDL-coated 6-well plates. DEM was diluted to corresponding molar dilution in DMEM/F12 supplemented with 15% FBS and penicillin/streptomycin (all from Thermo Fisher Scientific).

### Stereotactic surgery and LPS treatment

Mice at P21-28 were anesthetized and placed in the mouse stereotaxic frame (WPI) to secure the cranium. Then, the mice were injected with 250-500 nl of AAV (2.0 × 10^10^ GC/μl) at the rate of 100-200 nl/min into the medial prefrontal cortex (mPFC) by using a NanoFil syringe (WPI) and the following stereotactic coordinate; AP: +1.5 mm; ML: −0.2 mm; and DV: −1.8 mm from the bregma. Three to four weeks later, LPS (5 mg/kg) was injected intraperitoneally and the brains were harvested 48 h later.

### Immunohistochemistry

Mice were anesthetized and transcardially perfused with ice-cold PBS followed by 4% paraformaldehyde (PFA). Free floating sections (40 μm in thickness) were prepared by a Leica cryostat and then incubated in a blocking solution (PBS supplemented with 2% Normal Goat Serum, 1% BSA, 0.1% TritonX, 0.05% Tween-20, and 0.05% sodium azide) for 1 h at room temperature and then incubated at 4°C overnight with the following primary antibodies; rabbit anti-Iba-1 (019-19741, Wako), chicken anti-GFP (ab13970, Abcam), mouse anti-NeuN (MAB377, EMD Millipore), rabbit anti-S100β (ab4066, Abcam), goat anti-Iba1 (NB100-1028, Novus Biologicals), mouse anti-Oligo2 (MABN50, EMD Millipore), rabbit anti-GSTM1 (12412-1-AP, Protein Tech), and rabbit anti-GSTT2 (17622-1-AP, Protein Tech). After washing with PBS, the sections were further incubated with fluorophore-conjugated secondary antibodies for 2□h at room temperature, followed by DAPI staining for 10 min at room temperature. The sections were mounted on glass slides with Permafluor™ mounting medium (Thermo Fisher Scientific). Images were acquired with a Zeiss LSM510 confocal microscope and Zen software.

### Image analysis

For microglial analysis, Z-stack images were analyzed with Image J (NIH). Three images were taken from the mPFC of each mouse (n=3 mice per group). Microglia activation status was categorized as either “activated” or “resting” based on the morphology as previously described (53). Then, the percentage of “activated” microglia per total microglia was calculated per each image and averaged to represent the activation status of the corresponding mouse. Percentage of TNF-α expressing microglia per total microglia was also calculated in the mPFC in the same manner (n=3 mice per group).

### Enzyme linked immunosorbent assay (ELISA)

ELISA was performed using Ready-SET-Go!® Kits (eBioscience) for TNF-α, CCL2, and GM-CSF following the manufacturer’s protocol.

### Statistical analysis

Student’s t-test, one-way analysis of variance (ANOVA), and two-way ANOVA were used and *p* <0.05 was considered statistically significant. For multiple testing corrections, Tukey’s post-hoc test (for one-way ANOVA) and Sidak’s post-hoc test (for two-way ANOVA) were utilized.

Additional information can be found in **SI Materials and Methods.**

## Acknowledgements

We thank Minae Niwa, Sun-Hong Kim, Yian Chen, and Akiho Murata for technical help; Atsushi Kamiya for BV2 microglia cells; Z. J. Huang for AAV-LSL-GFP-shRNAmir constructs; and Fengyi Wan for advice. This work was supported by the grants of National Institutes of Health (MH093458 to S.K.; MH094268, MH105660, MH092443, and DA040127 to A.S.), RUSK (to A.S.), BBRF (to A.S.), and Stanley (to A.S.) foundations. S.K. was also supported by Johns Hopkins Medicine Discovery Fund and the Department of Psychiatry Venture Discovery Fund.

## References

1. Wang DD & Bordey A (2008) The astrocyte odyssey. Prog Neurobiol86(4):342–367.

2. Molofsky AV, et al.(2012) Astrocytes and disease: a neurodevelopmental perspective.Genes Dev26(9):891–907.

3. Clarke LE & Barres BA (2013)Emerging roles of astrocytes in neural circuit development. Nat Rev Neurosci14(5):311–321.

4. Khakh BS & Sofroniew MV (2015)Diversity of astrocyte functions and phenotypes in neural circuits. Nat Neurosci18(7):942–952.

5. Sofroniew MV (2014)Multiple roles for astrocytes as effectors of cytokines and inflammatory mediators. Neuroscientist20(2):160–172.

6. Ransohoff RM & Brown MA (2012)Innate immunity in the central nervous system. J Clin Invest122(4):1164–1171.

7. Meister A & Anderson ME (1983)Glutathione. Annu Rev Biochem52:711–760.

8. Grek CL, Zhang J, Manevich Y, Townsend DM, & Tew KD (2013)Causes and consequences of cysteine S-glutathionylation. J Biol Chem288(37):26497–26504.

9. Janssen-Heininger YM, et al.(2013) Emerging mechanisms of glutathione-dependent chemistry in biology and disease. J Cell Biochem114(9):1962–1968.

10. Kulak A, et al.(2013)Redox dysregulation in the pathophysiology of schizophrenia and bipolar disorder: insights from animal models. Antioxid Redox Signal18(12):1428–1443.

11. Butterfield DA (2015)Redox signaling in neurodegeneration. Neurobiol Dis84:1–3.

12. Miller AH, Haroon E, & Felger JC (2017)The Immunology of Behavior-Exploring the Role of the Immune System in Brain Health and Illness. Neuropsychopharmacology42(1):1–4.

13. Wraith DC & Nicholson LB (2012)The adaptive immune system in diseases of the central nervous system. J Clin Invest122(4):1172–1179.

14. Koga M, Serritella AV, Sawa A, & Sedlak TW (2016)Implications for reactive oxygen species in schizophrenia pathogenesis. Schizophr Res176(1):52–71.

15. Landek-Salgado MA, Faust TE, & Sawa A (2016)Molecular substrates of schizophrenia: homeostatic signaling to connectivity. Mol Psychiatry21(1):10–28.

16. Kano S, et al.(2013)Genome-wide profiling of multiple histone methylations in olfactory cells: further implications for cellular susceptibility to oxidative stress in schizophrenia. Mol Psychiatry18(7):740–742.

17. Emiliani FE, Sedlak TW, & Sawa A (2014)Oxidative stress and schizophrenia: recent breakthroughs from an old story. Curr Opin Psychiatry27(3):185–190.

18. Hayes JD, Flanagan JU, & Jowsey IR (2005)Glutathione transferases. Annu Rev Pharmacol Toxicol45:51–88.

19. Landi S (2000)Mammalian class theta GST and differential susceptibility to carcinogens: a review. Mutat Res463(3):247–283.

20. Petermann A, et al.(2009)GSTT 2, a phase II gene induced by apple polyphenols, protects colon epithelial cells against genotoxic damage. Mol Nutr Food Res53(10):1245–1253.

21. Beiswanger CM, et al.(1995)Developmental changes in the cellular distribution of glutathione and glutathione S-transferases in the murine nervous system. Neurotoxicology16(3):425–440.

22. Awasthi YC, Sharma R, & Singhal SS (1994)Human glutathione S-transferases. Int J Biochem26(3):295–308.

23. Al Nimer F, et al.(2013)Naturally occurring variation in the Glutathione-S-Transferase 4 gene determines neurodegeneration after traumatic brain injury. Antioxid Redox Signal18(7):784–794.

24. Sharma K, et al.(2015)Cell type- and brain region-resolved mouse brain proteome. Nat Neurosci18(12):1819–1831.

25. Tew KD & Townsend DM (2012)Glutathione-s-transferases as determinants of cell survival and death. Antioxid Redox Signal17(12):1728–1737.

26. Board PG & Menon D (2013)Glutathione transferases, regulators of cellular metabolism and physiology. Biochim Biophys Acta1830(5):3267–3288.

27. Townsend DM & Tew KD (2003)The role of glutathione-S-transferase in anti-cancer drug resistance. Oncogene22(47):7369–7375.

28. Jones JT, et al.(2016)Glutathione S-transferase pi modulates NF-kappaB activation and pro-inflammatory responses in lung epithelial cells. Redox Biol8:375–382.

29. Smeyne M, et al.(2007)GSTpi expression mediates dopaminergic neuron sensitivity in experimental parkinsonism. Proc Natl Acad Sci U S A104(6):1977–1982.

30. Rossignol DA, Genuis SJ, & Frye RE (2014)Environmental toxicants and autism spectrum disorders: a systematic review. Transl Psychiatry4:e360.

31. Kim SK, et al.(2015)Genetic Polymorphisms of Glutathione-Related Enzymes (GSTM 1, GSTT 1, and GSTP1) and Schizophrenia Risk: A Meta-Analysis. Int J Mol Sci 16(8):19602–19611.

32. Rodriguez-Santiago B, et al.(2010)Association of common copy number variants at the glutathione S-transferase genes and rare novel genomic changes with schizophrenia. Mol Psychiatry15(10):1023–1033.

33. Allen M, et al.(2012)Glutathione S-transferase omega genes in Alzheimer and Parkinson disease risk, age-at-diagnosis and brain gene expression: an association study with mechanistic implications. Mol Neurodegener7:13.

34. Lee YH, et al.(2015)Meta-analysis of associations between MTHFR and GST polymorphisms and susceptibility to multiple sclerosis. Neurol Sci36(11):2089–2096.

35. Zhang B, et al.(2013)Integrated systems approach identifies genetic nodes and networks in late-onset Alzheimer's disease. Cell153(3):707–720.

36. Burda JE & Sofroniew MV (2014)Reactive gliosis and the multicellular response to CNS damage and disease. Neuron81(2):229–248.

37. Liddelow SA, et al.(2017)Neurotoxic reactive astrocytes are induced by activated microglia. Nature541(7638):481–487.

38. Saijo K, et al.(2009)A Nurr1/CoREST pathway in microglia and astrocytes protects dopaminergic neurons from inflammation-induced death. Cell137(1):47–59.

39. Mayo L, et al.(2014)Regulation of astrocyte activation by glycolipids drives chronic CNS inflammation. Nat Med20(10):1147–1156.

40. Ponomarev ED, et al.(2007)GM-CSF production by autoreactive T cells is required for the activation of microglial cells and the onset of experimental autoimmune encephalomyelitis. J Immunol178(1):39–48.

41. Imai Y & Kohsaka S (2002) Intracellular signaling in M-CSF-induced microglia activation: role of Iba1. Glia40(2):164–174.

42. Hayden MS & Ghosh S (2008)Shared principles in NF-kappaB signaling. Cell132(3):344–362.

43. Ruland J (2011)Return to homeostasis: downregulation of NF-kappaB responses. Nat Immunol12(8):709–714.

44. Lou H & Kaplowitz N (2007)Glutathione depletion down-regulates tumor necrosis factor alpha-induced NF-kappaB activity via IkappaB kinase-dependent and -independent mechanisms. J Biol Chem282(40):29470–29481.

45. Aurnhammer C, et al.(2012)Universal real-time PCR for the detection and quantification of adeno-associated virus serotype 2-derived inverted terminal repeat sequences. Hum Gene Ther Methods23(1):18–28.

46. Kano S, et al.(2015)Clinical utility of neuronal cells directly converted from fibroblasts of patients for neuropsychiatric disorders: studies of lysosomal storage diseases and channelopathy. Curr Mol Med15(2):138–145.

47. Seshadri S, et al.(2015)Interneuronal DISC1 regulates NRG1-ErbB4 signalling and excitatory-inhibitory synapse formation in the mature cortex. Nat Commun6:10118.

48. Delenda C & Gaillard C (2005)Real-time quantitative PCR for the design of lentiviral vector analytical assays. Gene Ther12 Suppl 1:S36–50.

49. Chang K, Marran K, Valentine A, & Hannon GJ (2013)Creating an miR30-based shRNA vector. Cold Spring Harb Protoc2013(7):631–635.

50. de Vellis J & Cole R (2012)Preparation of mixed glial cultures from postnatal rat brain.Methods Mol Biol814:49–59.

51. Lee JK & Tansey MG (2013)Microglia isolation from adult mouse brain. Methods Mol Biol1041:17–23.

52. Ozeki Y, et al.(2011)A novel balanced chromosomal translocation found in subjects with schizophrenia and schizotypal personality disorder: altered l-serine level associated with disruption of PSAT1 gene expression. Neurosci Res69(2):154–160.

53. Thored P, et al.(2009)Long-term accumulation of microglia with proneurogenic phenotype concomitant with persistent neurogenesis in adult subventricular zone after stroke. Glia57(8):835–849.

